# Defence recognition of a stripe rust fungal effector is uncoupled from disease outcomes in wheat

**DOI:** 10.1101/2025.07.30.667774

**Authors:** Eric C. Pereira, Bayantes Dagvadorj, Rita Tam, Haoran Li, Danish Ilyas Baig, Mareike Möller, Miraclemario Raphael, Simon Williams, Sambasivam Periyannan, Florence Danila, John P. Rathjen, Benjamin Schwessinger

## Abstract

Plant resistance (*R*) and pathogen avirulence (*Avr*) gene interactions are central to pathogen recognition and disease resistance in crops. Functional characterisation of recognised *Avr* effectors of *Puccinia striiformis* f. sp. *tritici* (*Pst*) lags other key fungal pathogens of wheat. Here, we used a wheat protoplast-based screen to identify *Avr*/*R* interactions via the proxy of effector-induced defence responses in a set of diverse wheat cultivars. We identified an *Avr* candidate, termed *AvrPstB48,* that triggers defence responses in 16 out of 24 cultivars tested. *AvrPstB48* is hemizygous, and the *Pst* genome carries four divergent paralogs within a gene cluster. Analysis of these paralogs revealed partial redundancy in their ability to activate wheat defences and enabled us to identify a single amino acid in *AvrPstB4*8 that is necessary but not sufficient for defence activation. Notably, the activation of defence signalling by *AvrPstB48* in protoplasts did not directly correlate with disease outcomes. Whole-plant infection assays revealed that some cultivars which exhibited strong defence activation in the protoplast assay are susceptible to the *Pst* isolate Pst104E137A- from which *AvrPstB48* is derived. Comparison of infection dynamics of two wheat cultivars that differ in their *AvrPstB48* recognition capacity revealed a delay in disease progression in the recognising cultivar Avocet *S* compared to the non-recognising cultivar Morocco. While correlative only, our observations, combined with other recent reports, support a ‘recognize-then-suppress’ model of plant-pathogen interaction where disease outcomes are driven not only by simple *Avr/R* interactions but also by pathogen effectors that suppress defence signalling downstream of effector recognition.

## INTRODUCTION

Plant pathogenic fungi are major threats to crop yields globally. This is exemplified by wheat stripe (yellow) rust caused by *Puccinia striiformis* f. sp. *tritici* (*Pst*), which causes significant crop losses driven by the ongoing intercontinental spread and rapid host adaptation of this fungus (Hovmøller et al., 2010; 2025). On the molecular level, disease is caused by a large set of effector proteins that are secreted during wheat (Wang et al., 2023). These effectors modulate host cellular processes to facilitate host colonisation, including nutrient acquisition and suppression of defence responses (Zheng et al., 2023). The suppression of defence responses is especially important for biotrophic plant pathogens such as *Pst* which require living host tissue for prolonged periods to complete their lifecycles (Schwessinger, 2017). A subset of effectors designated avirulence (*Avr*) effectors are detected by host resistance (*R*) genes in certain genotypes and cultivars (Xia et al., 2020). *Avr* recognition triggers a myriad of defence responses that restrict pathogen colonisation, including transcriptional reprogramming and, at times, programmed cell death (Dodds & Rathjen, 2010). Several *Avr* genes from other wheat rust fungi have been cloned including *AvrSr13, AvrSr22, AvrSr27, AvrSr35, AvrSr50*, and *AvrSr62* from *P. graminis* f. sp. *tritici* (Arndell et al., 2024; Chen et al., 2017; Chen et al., 2024; Outram et al., 2024; Upadhyaya et al., 2021; Salcedo et al., 2017) and *AvrLr15* and *AvrLr21* from *P. triticina* (Cui et al., 2024; Shen et al., 2024). Similar advances have been reported for *Blumeria graminis*, with the identification of *AvrPm2* (Praz et al., 2017) and *AvrPm3* (Bourras et al., 2019), and *Zymoseptoria tritici* with the identification of *AvrStb6*, *Avr3D1* and *AvrStb9* (Amezrou et al., 2023; Meile et al., 2018; Zhong et al., 2017).

Disease outcomes are not always defined by *Avr/R* recognition events alone. Some effectors suppress defence signalling or modulate host metabolic pathways to subvert defence activation (Oliveira-Garcia et al., 2023; Ramírez-Zavaleta et al., 2022; Rhodes et al., 2022). Defence suppressive effectors can be categorised into causing effector-triggered susceptibility (ETS) or effector-triggered susceptibility II (ETS II), depending on the molecular targets and their sequential effect during plant infection (Jones & Dangl, 2006; Sohn et al., 2007; Win et al., 2012). ETS is defined as effector-suppression of the defence response downstream of the recognition of conserved pathogen molecules via pattern recognition receptors (Miya et al., 2007). ETS II is defined as effector-suppression of defence signalling downstream of *Avr/R* recognition events and can also be referred to as a ‘recognize-then-suppress’ model of plant-pathogen interactions.

Despite significant progress in cloning wheat stripe rust *R* genes, the identification of corresponding *Avr* genes in *Pst* has lagged far behind. To overcome these challenges, we adapted and optimised a rapid protoplast-based screening system for functional testing of *Pst* effector candidates (Wilson et al., 2024). We focused on hemizygous effectors identified in the fully phased genome assembly of the Australian *Pst* isolate Pst104E137A- (Tam et al., 2025). These effectors are of particular interest because hemizygosity implies that only a single allele of an effector needs to undergo a mutational event to avoid recognition in case of an avirulence effector. By screening these previously identified hemizygous effectors, we isolated a single avirulence effector candidate that is differentially recognised by a diverse set of wheat cultivars. This study characterises this effector termed *AvrPstB48* and its four paralogs using a protoplast screening system and diverse wheat cultivars. In addition to confirming their recognition, we analysed *AvrPstB48* sequence diversity, expression patterns during infection, recognition specificity and their potential impacts on disease outcome during infection.

## MATERIAL AND METHODS

### Candidate *Pst Avr* selection and gene synthesis

Hemizygous effectors were previously identified by analysing a time course of long-read transcriptomic datasets of Pst104E137A- as detailed in Tam et al. (2025). Briefly, transcripts were sampled from six conditions, each with four replicates, including ungerminated urediniospores and infected wheat leaves at 4, 6, 8, 10, and 12 days post-infection (dpi). All samples were subjected to direct Oxford Nanopore Technologies (ONT) cDNA sequencing. DESeq2 (Love et al., 2014) differential expression analysis was conducted on transcript read abundance to identify secretome genes upregulated (log_2_ fold change ≥ 2) early during infection (4 and 6 dpi) relative to dormancy. Inter-haplotype sequence analysis was then conducted to shortlist the candidates based on hemizygosity. The selected candidates were synthesised as gene blocks to facilitate precise assembly through Golden Gate cloning (Engler et al., 2014). Here, we describe the functional characterisation of one of these hemizygous effector candidates, ID Pst104E137_010232, referred to as *AvrPstB48*.

### Identification of *AvrPstB48* paralogs

The tblastn v2.16.0+ was used to search for *AvrPstB48* paralogs using its protein sequence against the *Pst*104E genome assembly (-evalue 1e-5) (Camacho et al., 2009). We applied a low identity cut-off of 0.4, considering effector proteins are typically sequence-divergent yet structurally conserved (Yu et al., 2023). The search results were used to locate the annotated gene hits. Multiple protein sequence alignments of *AvrPstB48* and the paralogs Pst104E137_025766 (*AvrPstB48_P1*), Pst104E137_025767 (*AvrPstB48_P2*) and Pst104E137_010231 (*AvrPstB48_P3*), were performed using MAFFT v7.490 (Katoh & Standley, 2013) and visually inspected in Geneious to confirm overall similarities in physicochemical properties of individual amino acids in the alignment.

### Effector prediction

We used full-length protein sequences of *AvrPstB48, AvrPstB48_P1, AvrPstB48_P2/P4*, and *AvrPstB48_P3* with SignalP6 to predict the presence and location of signal peptides (Teufel et al., 2022). We used respective protein sequences without signal peptide with EffectorP3 to predict the probability of the proteins to encode for effectors, including preference for cytoplasmic and/or apoplastic effector location (Sperschneider & Dodds, 2022).

### *In silico* structural modelling

The predicted tertiary structures of *AvrPstB48* and its paralogs were generated using the AlphaFold Server v3.0 (DeepMind, UK) (Abramson et al., 2024). Models were generated based on the protein sequences with the signal peptides removed. The predicted Template Modelling (pTM) score can be used to assess global model quality and domain orientation, where scores ≥0.5 are generally considered acceptable for reliable domain-level predictions (Jumper et al., 2021). The initial scores for all structural models were <0.5 (Table S1). However, we observed from those models that ∼50 residues at the N-terminus of each protein were unstructured. We also noted that the sequences of the proteins were rich in conserved cysteine residues. In other rust effectors, these have been shown to co-ordinate Zn²^+^ (Zhang et al., 2018; Outram et al., 2024). We therefore repeated the predictions with the N-terminal truncated and Zn²^+^ included. We observed an overall improvement in the pTM scores for all models >0.5 when the N-terminal region was removed, and the scores generally improved further when Zn was added to the prediction. The interface prediction score for Zn binding was, in most cases>0.7, indicating acceptable interaction/binding reliability (Abramson et al., 2024) (Table S1). The resulting models were visualised and analysed using PyMOL and ChimeraX v1.8 (UCSF) (Meng et al., 2023). Structural comparison of *AvrPstB48* and its paralogs *(P1, P2/P4*, and *P3*) was performed via superimposition in PyMOL, using *AvrPstB48* as the reference structure to identify conformational differences that may underlie functional divergence (Table S1).

### Golden Gate assembly of expression constructs

Synthetic double-stranded DNA fragments (gBlocks Gene Fragments; Integrated DNA Technologies (IDT), Coralville, IA, USA) encoding the hemizygous effector candidate *AvrPstB48*, its paralogs *AvrPstB48*_*P1*, *AvrPstB48*_*P2*, *AvrPstB48*_*P3*, and the mutants *AvrPstB48*_D133G and *AvrPstB48*_*P3*_G133D were synthesised with flanking *BsaI* recognition sites to generate an AATG overhang downstream of the long intergenic region and a GCTT overhang upstream of the short intergenic region to facilitate level 1 Golden Gate assemblies. The digested fragments were ligated into the pWDV1 expression vector, which had also been digested with *BsaI*. The reaction was cycled between 37°C for 3 minutes (to allow digestion) and 16°C for 4 minutes (for ligation) for 25 cycles, followed by a final incubation at 50°C for 5 minutes and 80°C for 5 minutes to inactivate the *BsaI* enzyme. Following the Golden Gate reaction, 5 µL of the reaction mixture was transformed into competent *Escherichia coli* cells. The cells were incubated on ice for 30 minutes, heat-shocked at 42°C for 30 seconds, then returned to ice for 5 minutes. SOC medium (200 µL; containing 2% tryptone, 0.5% yeast extract, 10 mM NaCl, 2.5 mM KCl, 10 mM MgCl₂, 10 mM MgSO₄, and 20 mM glucose) was added, and the mixture was incubated at 37°C for 1 hour with shaking at 250 rpm.

Transformed *E. coli* cells were plated on LB agar supplemented with 35 μg/mL chloramphenicol and incubated overnight at 37°C. Individual colonies were screened by colony PCR using gene-specific primers to confirm the presence of *Avr* candidate inserts (Table S2). PCR-positive clones were cultured in LB broth for plasmid extraction using a standard miniprep protocol (Qiagen). Extracted plasmids were subjected to restriction enzyme digestion with insert-specific restriction sites to validate correct plasmid architecture. Clones showing the expected digestion patterns were then selected for full-length plasmid verification via ONT sequencing. Whole-plasmid sequencing was performed using ONT MinION flow cells, and raw reads were analysed in Geneious Prime v2025.1.3 using Minimap alignment against the plasmid reference map to confirm correct insert orientation and base pair sequence integrity (Li, 2018) (Table S3).

### Plant materials and growth conditions for protoplast isolation

For each wheat cultivar (cv), Chinese 166, Avocet/*Yr1*, Heines VII, Nord Desprez, *Triticum spelta*, Avocet/*Yr5*, Heines Peko, Heines Kolben, Avocet/*Yr6,* Reichersberg 42, Avocet/*Yr7*, Compare, Avocet/*Yr8,* Clement, Avocet/*Yr9*, *Yr15*, Avocet/*Yr15*, Spalding prolific, Avocet/*YrSp*, Avocet S, Breakwell, Gabo, Coorong and Morocco, 15 seeds were sown in each pot (14 cm in diameter, 10 cm in depth, and approximately 1.5 L in volume). The plants were grown in controlled-environment growth cabinets at 18°C under a 16 h light/8 h dark photoperiod and a light intensity of 300 μmol/m²/s. Relative humidity was maintained at 60% during the light period and 70% during the dark period. Ten grams of soluble fertiliser (HORTICO) was dissolved in 4.5 L of water and applied to each tray of plants. Plants were watered every two days. At 8 days after sowing, leaves were harvested for protoplast isolation.

### Protoplast isolation

Wheat protoplasts were isolated using the protocol described by Wilson et al. (2023, 2024), with minor modifications. Briefly, protoplasts were isolated by making shallow incisions in the leaf epidermis of the first and second true leaf, followed by peeling the epidermis to reveal the underlying mesophyll cells. The excised leaf segments were incubated with the peeled surface in contact with a 0.6 M mannitol solution for 10 minutes. Subsequently, the segments were transferred to a Petri plate (4.5 cm) containing an enzyme solution composed of MES buffer (pH 5.7, 20 mM), mannitol (0.8 M), KCl (10 mM), cellulase (Yakult R-10) 1.5% w/v, macerozyme (Yakult R-10) 0.75% w/v, CaCl2 (10 mM), and BSA 0.1% w/v. The plate was wrapped in aluminium foil to prevent light exposure and placed on an orbital shaker set to 60 rpm for 3 h at room temperature. Following enzyme digestion, released protoplasts were filtered through a 40 μm nylon cell strainer. Protoplasts were then collected from the enzyme solution by gentle centrifugation at 100 g for 3 minutes using a 30 mL round-bottom tube, resuspended and washed in W5 solution (MES buffer, pH 5.7, 2 mM; KCl, 5 mM; CaCl_2_, 125 mM; NaCl, 154 mM) and incubated at 4°C for 40 minutes to allow settling., A 10 µl aliquot of the concentrated protoplasts was taken for cell quantification and quality assessment using a hemocytometer and light microscope. The protoplasts were left at 4°C for 40 minutes to allow settling. The W5 solution was then removed, and the protoplasts were resuspended to a final concentration of 300,000–350,000 cells/mL using MMG solution (MES buffer, pH 5.7, 4 mM; mannitol, 0.8 mM; MgCl_2_, 15 mM). The protoplasts were used immediately for subsequent transfection.

### Protoplast transfection

Plasmid DNA (1 μg/μL) was prepared using standard mini and maxi-prep protocols (SV Wizard kits, Promega, Madison, WI, USA). Protoplast isolation and PEG-mediated transfection followed the method described by Wilson et al. (2024), adapted to a 96-well plate format to increase scalability and throughput (Khan et al., 2022; Lloyd et al., 2022) with additional optimizations (Pereira et al., 2025).

All expression constructs were based on the self-replicating pWDV1 vector backbone and were driven by the maize ubiquitin (UBI) promoter (Table S3). This included (i) individual *Avr* effector candidates, (ii) wheat resistance (*R*) genes of positive control Sr50, and (iii) luciferase reporter constructs. Reporter activity was quantified using a dual-luciferase system adapted from Wilson et al. (2024), in which a defence-inducible reporter plasmid (UBI-pDefense) and a constitutive normalisation plasmid (UBI-pRedf) were co-transfected. These reporters use the same substrate but encode luciferases with distinct emission spectra, which were read using dedicated Lumi Green band-pass (∼505 – 560 nm) and Red NB long-pass (cut-on ∼600 nm) filters on a luminescence plate reader (Tecan, Mannedorf, Switzerland; Infinite 200Pro) to allow ratiometric normalisation of defence induction.

For each transfection, three classes of reactions were prepared in parallel: (i) *Avr* candidates under test, (ii) a positive control (*AvrSr50* + Sr50), and (iii) a negative control (*AvrSr50* alone, no matching *R* gene). In positive-control wells, 0.5 μg of the *AvrSr50* effector plasmid, 2 μg of the Sr50 resistance gene plasmid, and 5 μg of each reporter plasmid (UBI-pDefense and UBI-pRedf) were co-transfected per well. Negative-control wells received 0.5 μg of the *AvrSr50* plasmid plus 5 μg of each reporter plasmid but no *Sr50* plasmid, to confirm that defence reporter activation required a cognate *R* gene. *Avr*-only reactions for candidate effectors were assembled analogously, using 1 μg of the *Avr* candidate plasmid and 5 μg of each reporter plasmid, without an *R* gene. Plasmid DNA mixes were dispensed into sterile 96-well V-bottom plates (Costar). Freshly isolated wheat protoplasts (50 μL per well) were added, followed by 50 μL of PEG solution (40% w/v PEG 4000, 0.2 M mannitol, 100 mM CaCl₂). The suspension was gently mixed by pipetting up and down five times, then incubated for 10 min at room temperature to allow DNA uptake. Transfection was stopped by adding 200 μL of W5 solution (154 mM NaCl, 125 mM CaCl₂, 5 mM KCl, 2 mM MES, pH 5.7). The plates were then incubated under low-light conditions at room temperature for 16 h to permit expression prior to luminescence quantification.

### Monitoring luciferase expression

After the 16h incubation, protoplasts were prepared for luminescence measurement. The W5 supernatant was removed gently using a multichannel pipette without disturbing the settled protoplast pellet. Fifty μL of 1× cell lysis buffer (#E3971; Promega) was added, and the samples were gently mixed by pipetting 5 times, then incubated for 15 minutes to ensure complete lysis. Fifty μL of lysate was collected and transferred to an opaque, flat-bottom, white, solid polystyrene 96-well plate (Corning 96-well NBS Microplate, #CLS3600). The total luminescence was measured using a plate reader (Tecan, Mannedorf, Switzerland; Infinite 200Pro, 1000 ms integration, 0 ms settle) without adding substrate to measure background signal. Immediately after this baseline read, 50 μL of Steady-Glo luciferase substrate (Promega, #E2520) was added to each well, and the luminescence was measured again. The plate reader was programmed first to measure total luminescence without filters, using an integration time of 1000 ms and a settle time of 0 ms. Subsequently, luminescence was recorded sequentially through the installed Lumi Green band-pass (∼505–560 nm) and Red NB long-pass (cut-on ∼600 nm) filters under the same acquisition settings. These filtered measurements allowed separation of the defence-inducible reporter (UBI-pDefense) and the constitutive normalisation reporter (UBI-pRedf), which use the same substrate but emit at different wavelengths.

### Protoplast RT-PCR

Wheat protoplasts were prepared as described above and dispensed at 900 µL per replicate (3.5 × 10⁵ cells mL⁻¹ in MMG buffer) for transfection. For each transfection, 9 µg of the candidate expression plasmid was co-transfected with 5 μg of each reporter. Transfections were performed in four independent biological replicates. After 16 h at room temperature under low light, 50 µL of each suspension was removed for luciferase quantification, and the remaining protoplasts were pelleted by centrifugation at 300 x g for 3 min. The supernatant was removed, snap-frozen in liquid nitrogen, and stored at −80 °C for RNA extraction.

Total RNA was isolated with the Direct-zol RNA MiniPrep Plus kit (Zymo Research) following the manufacturer’s instructions, including on-column DNase I treatment. RNA integrity was verified by agarose gel electrophoresis, and concentration and purity were measured with a Qubit fluorometer and a NanoDrop spectrophotometer. For cDNA synthesis, 1 µg of total RNA was reverse-transcribed using the High-Capacity cDNA Reverse Transcription Kit (Thermo Fisher Scientific) according to the manufacturer’s protocol.

Gene-specific primers targeting *AvrPstB48* and the paralog *AvrPstB48*_*P3* were designed in Geneious. Wheat 18S rRNA (Ta-18S) primers served as an endogenous reference control (Table S2).

Conventional endpoint RT-PCR was performed with GoTaq DNA polymerase (Promega) in 10 µL reactions with 3 µL cDNA template. Thermal cycling used a standard program (1 cycle of 95 °C for 1 min and 30 s; 30 cycles of 95 °C for 30 s, primer-specific annealing temperature of 50 °C for 30 s, and 72 °C for 2 min; final extension 72 °C for 5 min). Empty-vector (EV) transfections were included as negative biological controls; no-template controls (NTC) were run for each primer pair. Amplicons (5 µL per reaction) were resolved on 1% agarose in 1× TAE, stained, and imaged alongside a 1 kb Plus DNA Ladder (NEB #N3200).

### Inoculation of wheat plants with *Pst*

Fresh spores of *Pst* isolate Pst104E137A- (Schwessinger et al., 2018) were collected from infected leaf tissue. Spores were suspended in 3M Novec solution and measured using a CellDrop Automated Cell Counter (DeNovix Inc.) to a concentration of approximately 5×10^5^ spores/mL. Using a paintbrush, spores were inoculated on the upper surface of the first-emerged leaf of wheat cv Avocet *S* and Morocco 14 days after sowing at the same conditions described above. Infected plants were lightly sprayed with clean deionised water before being placed in an opaque, 25-L plastic container (33 cm x 22.5 cm x 47 cm) with a lid. Containers with infected plants were wrapped with black plastic bags and incubated at 10°C in the dark to initiate *Pst* germination. After 48 hours, the infected plants were removed from the container and transferred back to the growth cabinet. Each infected leaf was assessed according to Dracatos et al. (2016) over 10 or more days post-infection (dpi).

### Hyphae staining of infected wheat leaves

Staining was performed as described by Redkar *et al*. (2018). Freshly harvested leaf tissues were placed in 2 mL tubes containing 100% ethanol for three days or until the majority of the chlorophyll had been removed. When the tissues were nearly devoid of colour, ethanol was replaced with 10% aqueous potassium hydroxide and incubated on a dry heating block at 85°C for two hours. Cleared tissues were carefully washed 10 times in 1x phosphate-buffered saline (PBS Thermo Fisher Scientific, 28237), pH 7.4. Propidium iodide (PI; Sigma-Aldrich, P4170) was mixed with wheat germ agglutinin conjugated with AlexaFluor488 (WGA-AF488; Thermo Fisher Scientific, W11261) to create a PI-WGA AF488 staining solution (20 μg/mL PI, 10 μg/mL WGA-AF488, and 0.1% Tween 20 in 1× PBS, pH 7.4). After discarding the last PBS wash, PI-WGA-AF488 staining solution was added to the 2 mL tubes to fully submerge the tissues. Vacuum infiltration was applied at 7 millibars, repeated 5 times for 2 minutes each, with 2-minute vacuum release in between. Stained tissues were washed twice in 1x PBS to remove excess staining solution, transferred to fresh 1x PBS and stored at 4°C in the dark.

### Callose staining of infected wheat leaves

Freshly harvested leaf tissues were placed in 2 mL tubes containing 1x PBS, pH 7.4 and vacuum infiltrated at 7 millibars until they sank to the bottom of the tube. Prior to staining, leaf tissues were hand-sectioned paradermally using a razor blade. Eight pieces of sectioned leaf tissue were placed in each well of the 96-well plate and washed with 200 μL 1x PBS. Tissues were stained with 0.1% aniline blue (AB; Sigma-Aldrich, 415049) in 1x PBS for 60 mins with vacuum infiltration and washed twice in 1x PBS. WGA-AF488 staining solution (10 μg/L WGA-AF488 and 0.1% Tween 20 in 1× PBS, pH 7.4) was added to the samples and vacuum infiltrated at 7 millibars for 5 times, 2 minutes each, with 2-minute vacuum release in between. Samples were washed twice in 1x PBS, transferred to fresh 1x PBS and stored at 4°C in the dark.

### Confocal fluorescence microscopy

Stained leaf tissues were mounted onto a glass slide with 1x PBS and imaged under a TCS SP8 confocal microscope (Leica Microsystems) using 20x (PI-WGA-AF488) and 63x (AB-WGA-AF488) water-immersion objectives. WGA-AF488-stained fungal structures were visualised at 500-540 nm following excitation with an Argon laser at 488 nm. PI-stained cell wall was visualised at 580-630 nm following excitation with a DPSS laser at 561 nm. AB-stained callose was visualised at 430-500 nm following UV excitation at 405 nm. To obtain a composite image with greater depth of field, Z-stacking was performed with each optical section captured at a 0.1 μm interval. Images were processed (e.g., maximum projection, brightness and contrast) using Leica Application Suite X software (LASX, Leica Microsystems).

### Data collection and statistical analysis

Differences in normalised luminescence values were assessed using one-way analysis of variance (ANOVA) followed by Tukey’s Honestly Significant Difference (HSD) post hoc test (Wilson et al., 2024). The infectious hyphae cover area per leaf surface area was quantified using images obtained from PI-WGA-AF488-stained leaf tissues. Meanwhile, the callose signal area was quantified using images obtained from AB-WGA-AF488-stained leaf tissues. These quantifications were performed using ImageJ software (National Institutes of Health) on a Wacom Cintiq graphics tablet (Wacom Technology). Statistical analysis of data obtained from confocal images was performed using Welch’s t-test.

## RESULTS

### A hemizygous stripe rust candidate effector triggers defence signalling in wheat protoplasts

The hemizygous effector candidates were identified previously from the nuclear haplotype-phased genome of the *Pst* reference isolate Pst104E137A- (Tam et al., 2025). Here we describe the functional characterisation of one of these hemizygous effector candidates referred to here as *AvrPstB48*. The name *AvrPstB48* reflects its predicted avirulence function “Avr” in *Pst*, with “B” representing its broad recognition pattern across diverse wheat cultivars, and “48” corresponding to its internal candidate ID in the screening pipeline. Additional hemizygous effector candidates, including *Pst104E137_008104* (#47) and *Pst104E137_024713* (#49), were selected using the same criteria and served as comparisons for validation assays (Tam et al., 2025).

Because *AvrPstB48* expression pattern aligned with a function during plant infection (Tam et al., 2025), we cross-validated its role as effector using the machine learning tool EffectorP3 (Sperschneider & Dodds, 2022). *AvrPstB48* was predicted as an effector with a most likely intracellular localisation. We therefore decided to perform transient expression of *AvrPstB48* in wheat protoplasts with its signal peptide removed to allow for intracellular localisation. Using a dual luciferase-based defence reporter assay with the D14 defence promoter (Wilson et al., 2024), we observed a robust and specific increase in ratiometric luminescence following co-expression of *AvrPstB48* and the defence reporter, indicating a recognition-dependent defence response in Avocet *Yr5* and Avocet *S* (Figure 1). The increase in ratiometric luminescence observed when expressing *AvrPstB48* was similar to the *AvrSr50/Sr50* gene pair, which served as a positive control to validate the assay performance (Wilson et al., 2024). By contrast, two other hemizygous effector candidates, #47 and #49, did not induce measurable reporter activity being similar to the negative control, *AvrSr50* (Figure 1). Together, these results are consistent with *AvrPstB48* encoding a recognised effector activating host defence responses in specific wheat cultivars (Table S5).

**Figure 1.**
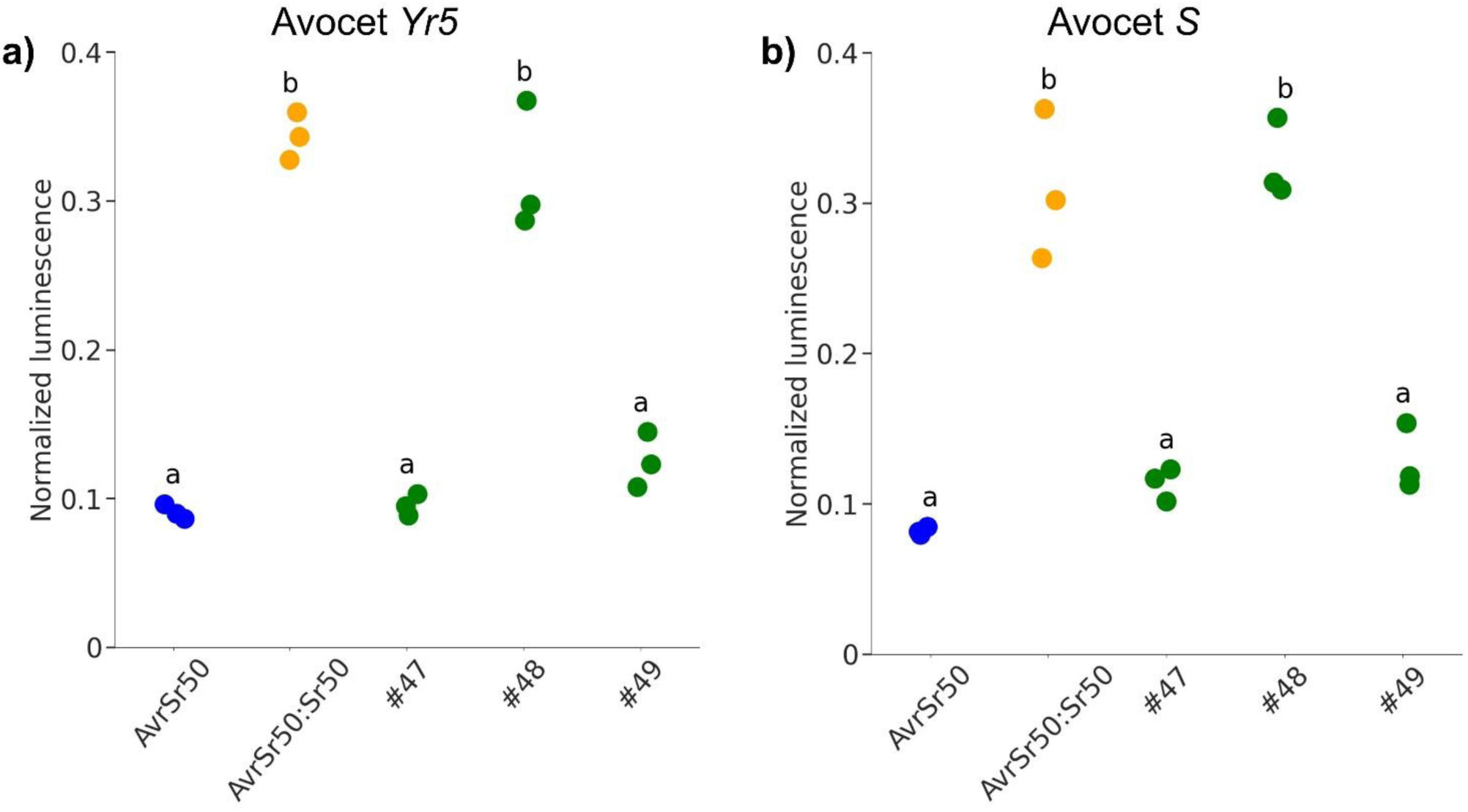
*pDefense* D14 promoter activation triggered by *AvrPstB48* in Avocet *Yr5* and Avocet *S* protoplasts. Dual luciferase-based reporter assays were performed using wheat protoplasts from wheat cultivars Avocet *Yr5* **a)** and Avocet *S* **b)** to assess defence signalling. The x-axis shows the plasmid combinations tested, including candidate effectors *Pst104E137_008104* (#47), *Pst104E137_010232* (*AvrPstB48*), and *Pst104E137_024713* (#49), along with the positive control pair *AvrSr50/Sr50*. The y-axis represents normalised luminescence values, measured as the ratio of luciferase activity driven by the D14 promoter to the internal control UBI-pRed. *AvrPstB48* triggered a significant increase in reporter activity, indicating activation of defence responses. Statistical differences were analysed using one-way ANOVA followed by Tukey’s Honestly Significant Difference (HSD) test; Conditions with different letters are significantly different from each other (*p* < 0.05).

Given the strong defence promoter activation observed for *AvrPstB48*, further analysis was conducted to identify and characterise its paralogs. Sequence homology analysis was performed by tblastn and revealed two divergent paralogs located in close proximity on each haplotype of chromosome 10, *AvrPstB48_*P1, *AvrPstB48_*P2, *AvrPstB48_P3* and *AvrPstB48_*P4. Among these allelic pairs, *AvrPstB48_*P2 and *AvrPstB48_*P4 share 100% nucleotide identity, whereas *AvrPstB48_*P1 and *AvrPstB48_P3* exhibit 98.2% nucleotide identity. Relative to AvrPstB48, the *P2/P4* pair is more closely related (90.8%), whereas the *P1/P3* pair is more divergent (57.6% identity to *AvrPstB48_P1*) (Figure 2a). Consistent with this, protein sequence alignments show that AvrPstB48_P2/P4 are highly similar to AvrPstB48, whereas AvrPstB48_P1 and AvrPstB48_P3 carry multiple non-synonymous substitutions and are more divergent at the amino-acid level (Figure 2b). AlphaFold3 modelling supported a shared fold among AvrPstB48, P1, P2/4, and P3. Backbone superposition yielded pairwise RMSDs of ∼0.5–1.5 Å, with divergence concentrated in the C-terminal helical region and adjacent loops (Fig. 2c). Global pTM values (≈0.50–0.57) indicate these are hypothesis-generating rather than definitive models; nevertheless, they are consistent with a conserved core and local surface variation between the paralogs. The locations of these variations correspond to clusters of substitutions in the multiple sequence alignment (Fig. 2b) and may modulate protein stability or interactions with host targets. Temporal expression profiling across infection stages revealed distinct expression patterns among the paralogs (Figure 2d). *AvrPstB48* exhibited peak expression at 8 days post-inoculation (dpi), reaching its highest levels relative to the other paralogs. All paralogs also peaked at 8 dpi, although expression intensity varied, with *P1*, *P2/P4*, and *P3* remaining moderately expressed throughout the infection time course. None of the paralogs were expressed in ungerminated spores, supporting their likely function in host-pathogen interactions.

**Figure 2.**
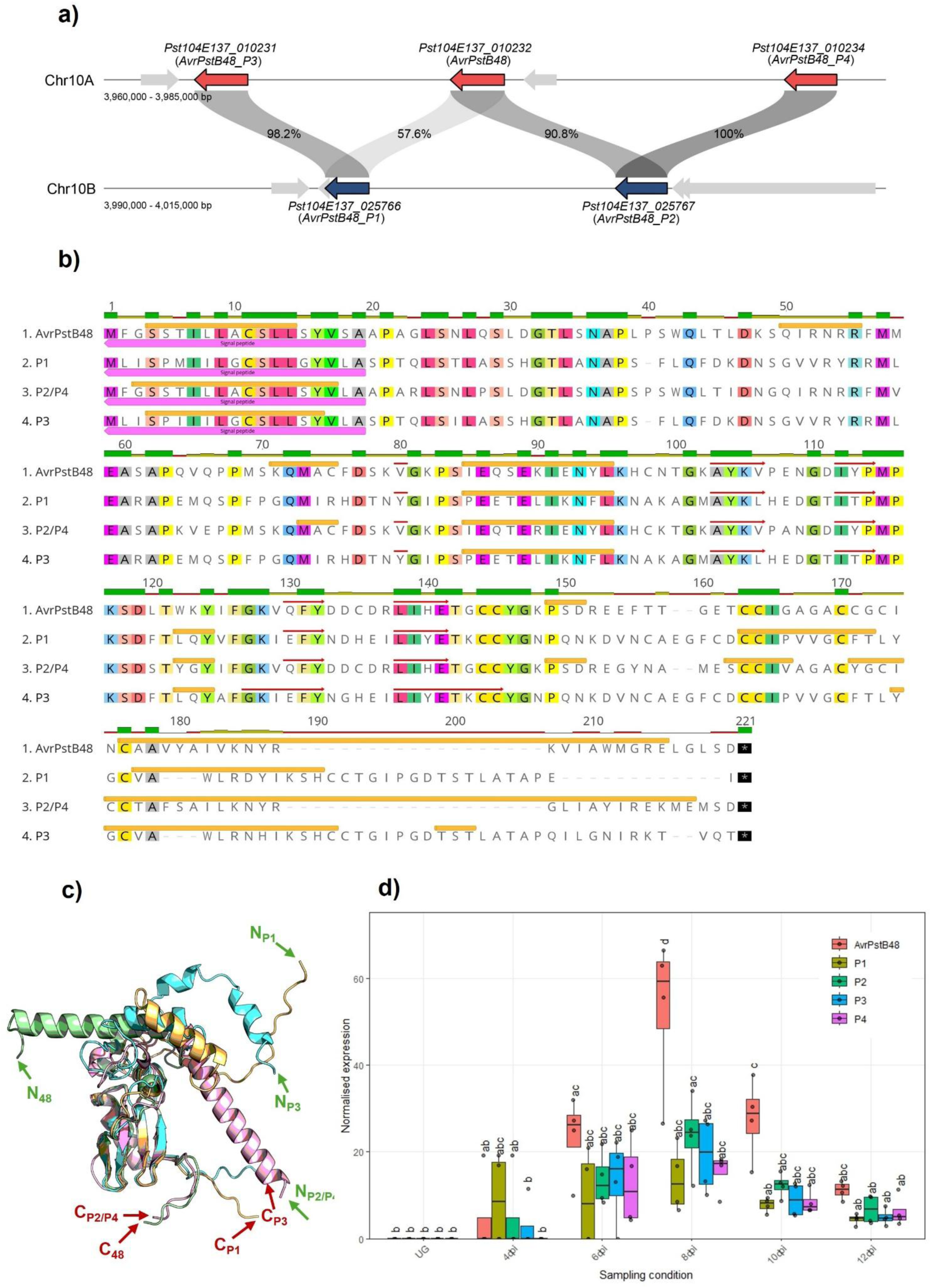
Genomic, evolutionary, and structural characterisation of the candidate effector gene *AvrPstB48* and its paralogs. **a)** Genomic organisation and synteny of *AvrPstB48* (*Pst104E137_010232*) and its paralogs (P1: *Pst104E137_025766*; P2: *Pst104E137_025767*; P3: *Pst104E137_010231*; P4: *Pst104E137_010234*) across homologous regions of Chromosomes 10A and 10B, with genes shown as red or blue arrows, respectively. Grey-shaded blocks and contained numbers represent pairwise nucleotide sequence identity (%) between all paralogs. **b)** Multiple sequence alignment of *AvrPstB48* and its paralogs, showing conservation at the amino acid level. Identical or similar residues are highlighted using the Clustal X colour scheme, which assigns colours based on amino acid conservation and properties. Conserved motifs are evident across the sequences. Dashes indicate gaps in the alignment. Pink shading marks the predicted N-terminal signal peptide; orange bars denote predicted α-helical segments; red arrows denote predicted β-strands mapped from the AlphaFold3 structural model onto the multiple-sequence alignment of *AvrPstB48* and its paralogs (*P1, P2/P4, P3*). **c)** Superposition of AlphaFold3 models for *AvrPstB48* and its paralogs. Global backbone similarity is high (pairwise RMSD ∼0.5–1.5 Å), with most divergence at the C-terminal helical region. Predicted model confidences (pTM ∼0.50–0.57) indicate hypothesis-generating structures rather than definitive models. Colour key: *AvrPstB48* (green), *P1* (orange), *P2/4* (pink), *P3* (light blue). The N-terminus are represented by green arrows and the C-terminus by red arrows, and includes *AvrPstB48* (C48) paralog (PX) labelling. **d)** Expression profile of *AvrPstB48* and its paralogs in ungerminated urediniospores (UG) and infected wheat leaves at 4, 6, 8, 10 and 12 days post-infection (dpi) *in planta*. RNA-seq abundance was normalised using DESeq’s median of ratio method and plotted as box plots with four biological replicates. Analysis of variance (ANOVA) was performed using a two-way ANOVA with sampling condition (UG, 4, 6, 8, 10 and 12 dpi) and effector identity (*AvrPstB48, P1, P2, P3* and *P4*) as fixed factors, followed by post hoc Tukey’s Honestly Significant Difference (HSD) test. Sampling condition –effector combinations marked with distinct letters indicate that they are significantly different (*p* < 0.05) according to Tukey’s post hoc multiple-comparisons test. Boxes represent the interquartile range with the horizontal line inside the box indicating the median.

### *AvrPstB48* recognition is conserved across multiple wheat cultivars

To further assess defence promoter activation triggered by *AvrPstB48*, we tested their activities in additional wheat cultivars including Avocet S, Morocco, *Triticum spelta Yr5*, Chinese 166 and Gabo, using three previously validated defence reporter promoters (D2, D14, and D15) (Wilson et al., 2024). Significant activation of each promoter was observed in response to *AvrPstB48* expression in Avocet *S* (Figure 3a), *T. spelta Yr5* and Gabo (Figures S1a, S1b), indicating effector recognition. In contrast, no increase in ratiometric luminescence signal was detected in Morocco (Figure 3b) or Chinese 166 (Figure S1c), suggesting a lack of *AvrPstB48* recognition in these cultivars. Additional screening of 16 wheat cultivars confirmed consistent activation in all near-isogenic Avocet lines, while eight cultivars showed no measurable response (Table S5). Importantly, the positive control (*AvrSr50/Sr50*) consistently induced strong luminescence across all tested backgrounds, validating assay performance.

**Figure 3.**
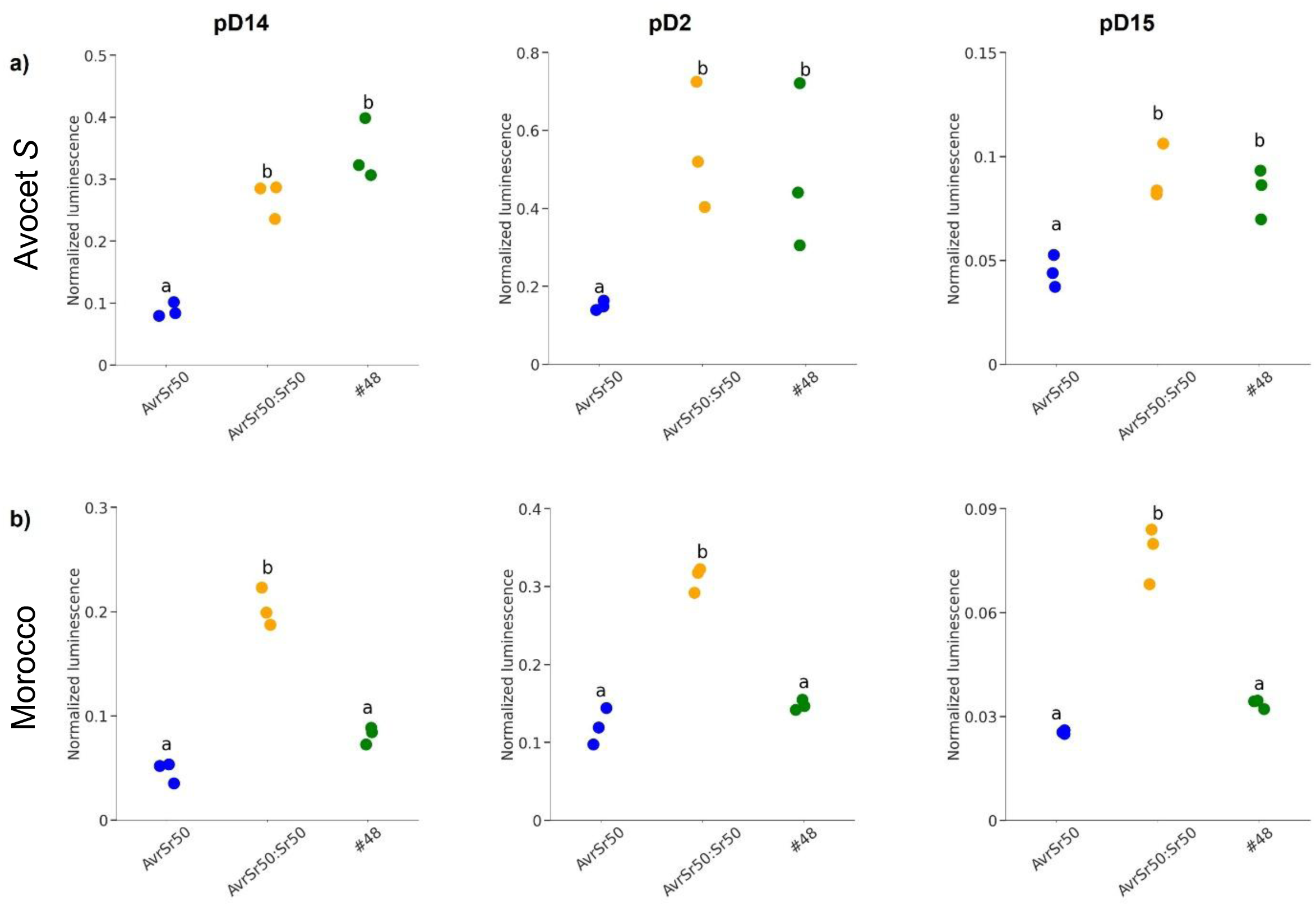
Differential activation of defence reporters by *AvrPstB48* reveals genotype-dependent recognition. Normalised luminescence signals from dual luciferase-based reporter assays using three promoter constructs (pD14, pD2, pD15) and UBI-pRedf in **a)** Avocet S and **b)** Morocco. *AvrSr50* and *AvrSr50:Sr50* were used as negative and positive controls, respectively. One-way analysis of variance (ANOVA) followed by Tukey’s Honestly Significant Difference (HSD) test was used to compare treatments. Conditions with different letters are significantly different from each other (*p* < 0.05).

### *AvrPstB48* paralogs *AvrPstB48_P1* and *AvrPstB48_P2/P4,* but not *AvrPstB48_P3,* are recognized in multiple wheat cultivars

To assess the functional redundancy of *AvrPstB48*, we tested recognition of its paralogs in the same set of wheat cultivars, Avocet S, Morocco, *T. spelta Yr5*, Gabo, and Chinese 166, using activation of the D14 promoter as a readout. Defence activation was detected for *AvrPstB48_*P1 and *AvrPstB48_*P2/P4 in Avocet *S* (Figure 4a), *T. spelta Yr5,* and Gabo (Figures S2), suggesting a similar recognition spectrum to *AvrPstB48*. *AvrPstB48_P3* failed to activate the D14 defence promoter in any of the tested cultivars. No induction was observed for any of the paralogs tested in Morocco (Figure 4b) and Chinese 166 (Figure S2).

**Figure 4.**
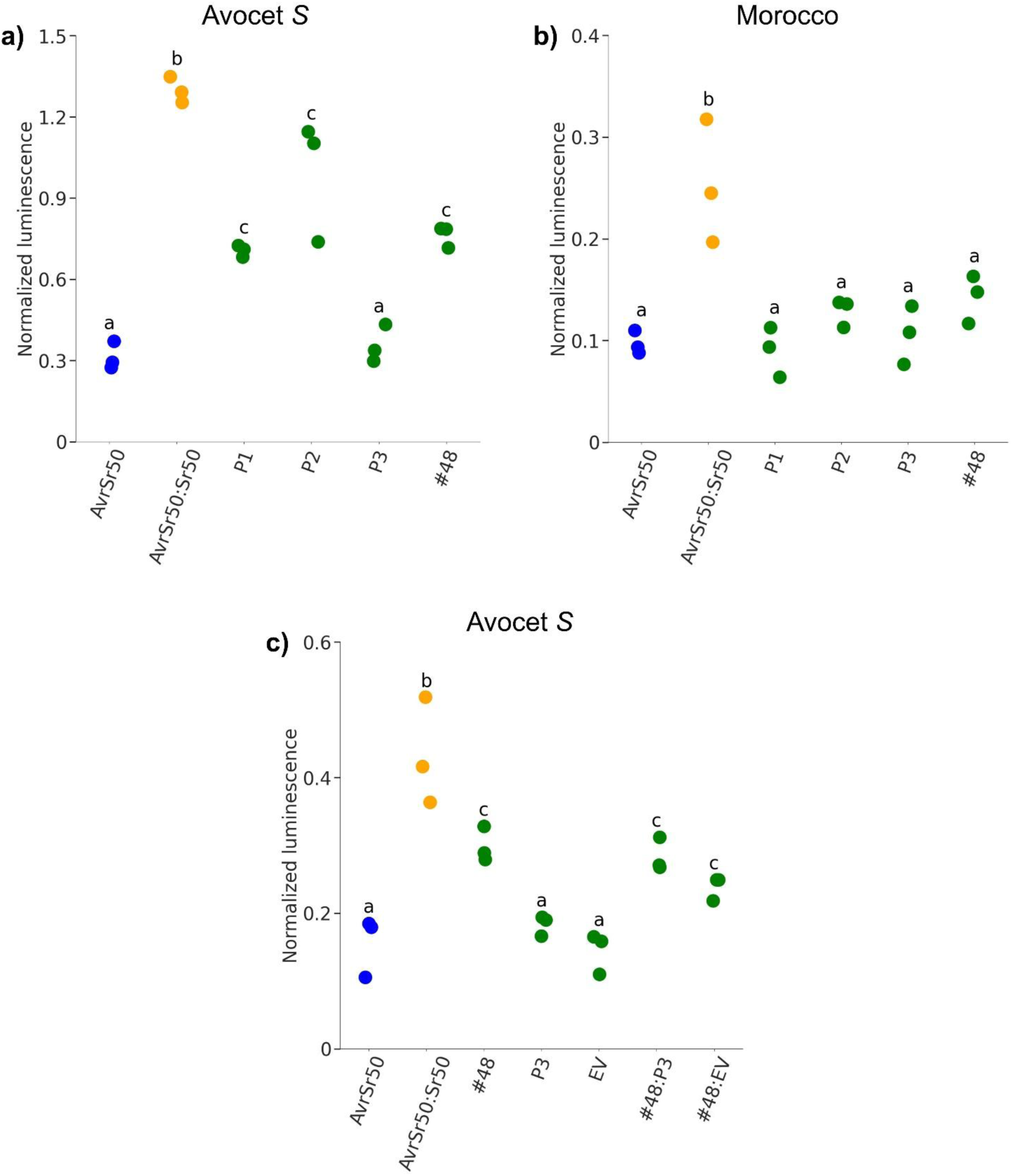
Differential activation of defence reporters by *AvrPstB48* paralogs reveals genotype-dependent recognition. Normalised luminescence signals from dual luciferase-based reporter assays using the pD14 promoter construct and UBI-pRedf in **a)** Avocet S, **b)** Morocco and **c)** Co-expression of *AvrPstB48* and *AvrPstB48_P3* (#48:P3) and *AvrPstB48* and Empty Vector (#48:ev) at a 1:1 plasmid ratio in Avocet S. *AvrSr50* and *AvrSr50:Sr50* were included as negative and positive controls. One-way analysis of variance (ANOVA) followed by Tukey’s Honestly Significant Difference (HSD) test was used to compare treatments. Conditions with different letters are significantly different from each other (*p* < 0.05).

Given that *AvrPstB48_P3* shares overall structural and sequence similarity with *AvrPstB48* (Figure 2), we investigated whether it could interfere with *AvrPstB48*-triggered defence signalling. In co-expression assays in Avocet *S* protoplasts, *AvrPstB48* was delivered either alone or together with *AvrPstB48*_P3 at a 1:1 plasmid ratio. Co-expression did not reduce relative ratiometric luminescence signal induction when compared to *AvrPstB48* alone, indicating that *AvrPstB48*_*P3* does not act as a competitive suppressive paralog when co-expressed in wheat protoplasts (Figure 4c). To verify effector transcript accumulation for both *AvrPstB48* and *AvrPstB48*_*P3*, we performed endpoint RT-PCR on cDNA from protoplasts transfected with the corresponding expression constructs. Gene-specific primer pairs yielded the expected amplicons for *AvrPstB48* and *AvrPstB48*_*P3* only in the matched transfection. No bands were detected in empty vector (EV) controls amplified with either primer set. The wheat 18S rRNA control amplified in all samples, supporting cDNA quality (Figure S3).

### Aspartic acid at position 134 is necessary but not sufficient for defence activation

To investigate why *AvrPstB48*_*P3* does not trigger defence promoter activation in Avocet S, we explored potential structural and biochemical differences among *AvrPstB48* and its paralogs. Structural predictions using AlphaFold3 revealed that all paralogs adopt a broadly conserved fold (Figure 2d). Closer inspection revealed several amino acid substitutions specific to *AvrPstB48_P3*, particularly in surface-exposed loops that may be involved in effector recognition and receptor interactions. One notable difference occurs at position 134, where *AvrPstB48, AvrPstB48_P1*, and *AvrPstB48_P2* encode an aspartic acid (D), while *AvrPstB48_P3* encodes a glycine (G) (Figure 5a). This substitution replaces a negatively charged residue (D) with a small, nonpolar amino acid (G) at a position that is predicted to be surface-exposed on a protruding loop. This makes the D134G substitution a prime candidate for functional testing.

**Figure 5.**
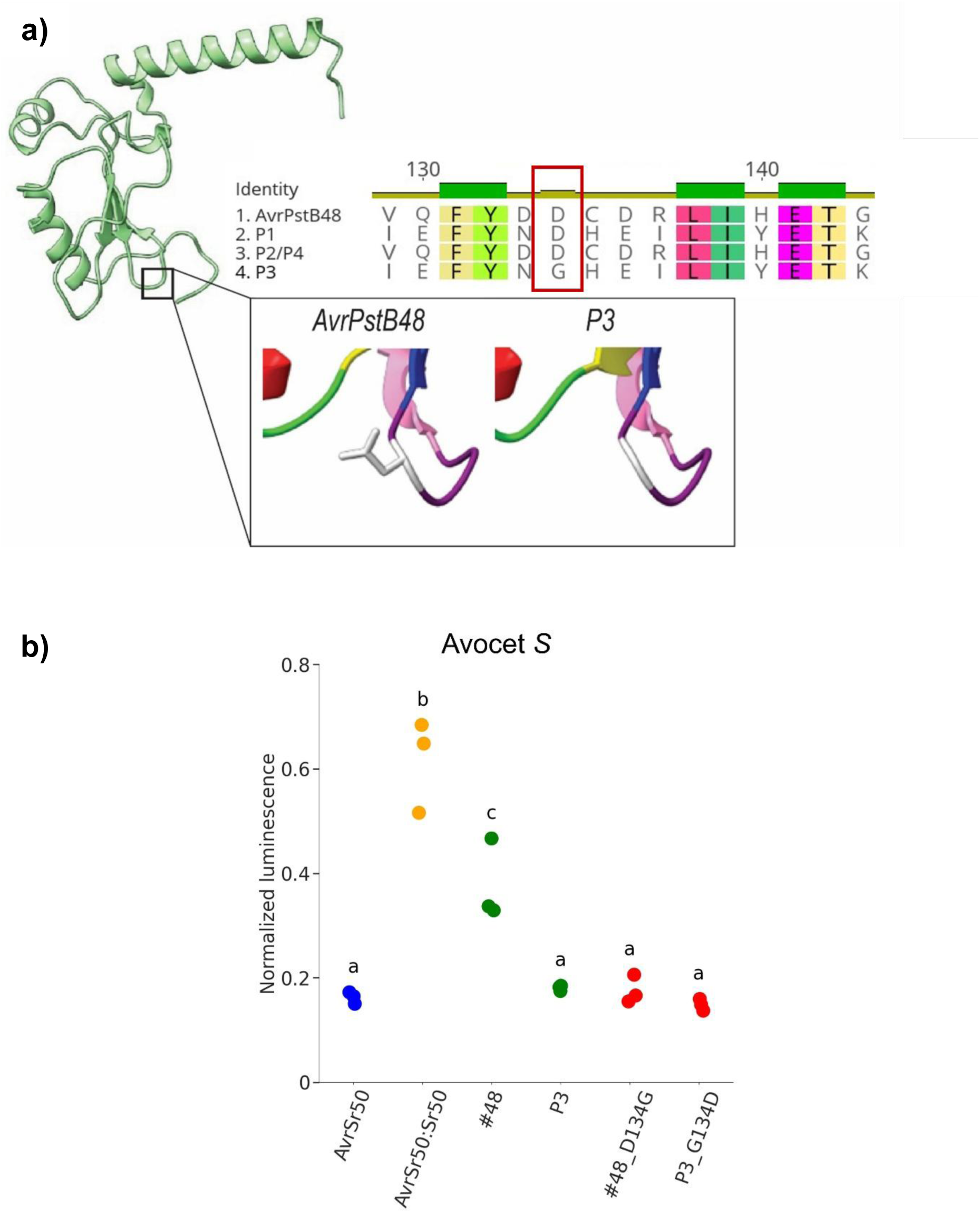
A single amino acid (D134) is necessary but not sufficient for recognition of *AvrPstB48* and its paralog *AvrPstB48_P3*. **a)** Predicted structures of *AvrPstB48* and *AvrPstB48*_P3 were generated using AlphaFold3. The surface-exposed amino acid position 134 is highlighted where *AvrPstB48*, *P1*, and *P2/P4* encode an aspartic acid (D), while P3 encodes a glycine (G). **b)** Normalised luminescence signals from dual luciferase-based reporter assays using the pD14 promoter and UBI-pRedf. One-way analysis of variance (ANOVA) followed by the Tukey Honestly Significant Difference (HSD) test was used to compare the treatments. Conditions with different letters are significantly different from each other (*p* < 0.05).

To further investigate the functional significance of the amino acid variation at position 134, synthetic gene fragments were designed to encode *AvrPstB48* carrying an aspartic acid to glycine substitution (D134G), and *AvrPstB48_P3* carrying the reverse substitution (G134D). These constructs were tested for their ability to induce defence promoter activation in Avocet *S* using the D14 promoter-luciferase reporter system in wheat protoplasts. AvrPstB48_D134G failed to activate the defence reporter when compared to the increase of ratiometric luminescence when expressing wild-type *AvrPstB48* (Figure 5b). This indicates that the D134G substitution is sufficient to disrupt recognition. The reciprocal mutant AvrPstB48_P3_G134D also failed to induce defence promoter activity, suggesting that the aspartic acid at position 134 is not the only residue contributing to the avoidance of recognition in *AvrPstB48_P3*.

### *AvrPstB48*-triggered reporter activation in protoplasts does not correlate fully with *Pst* resistance *in planta*

We assessed infection outcomes using the Australian pathotype Pst104E137A-(Schwessinger et al., 2018) to evaluate whether the *AvrPstB48*-triggered reporter activation observed in wheat protoplasts corresponded to resistance *in planta*. Wheat lines previously tested in the protoplast assay were inoculated and disease severity scored at 10 days post-inoculation (dpi) using a 0–9 infection type (IT) scale (Hovmøller et al., 2017). Reporter activation in protoplasts did not correlate with disease outcomes in all cultivars tested. While *T. spelta Yr5* and Chinese 166 exhibited full resistance (IT = 0), Gabo, Morocco, and Avocet *S* were highly susceptible (IT = 9), despite *AvrPstB48* triggering defence promoter activation in Gabo and Avocet *S* (Figure S4).

### *AvrPstB48* recognition correlates with delayed infection progression in the susceptible wheat cultivar Avocet S

To explore whether *AvrPstB48* recognition might influence infection dynamics even in a susceptible genotype, we compared Avocet *S* and Morocco. Both of these are susceptible to the *Pst* isolate Pst104E137A-, but Avocet *S* responds to *AvrPstB48* expression in the protoplast assay, whereas Morocco does not. At 8 dpi, neither line exhibited macroscopic ureodiniospore formation. By 10 dpi, both displayed visible rust symptoms, however, Morocco developed more extensive ureodiniospore coverage, indicating a higher sporulation capacity and greater disease severity (Figure 6A). Confocal microscopy supported these observations, revealing more abundant and earlier hyphal growth in Morocco compared to Avocet *S* at 8 dpi (Figure 6B). Although both lines were heavily colonised by 10 dpi, Avocet *S* retained localised areas with reduced fungal presence, consistent with greater disease severity. Quantification of infectious hyphal cover over time showed broadly similar colonisation dynamics between the two genotypes, with Morocco tending to exhibit slightly higher mean hyphal coverage at later time points (Figure 6D).

**Figure 6.**
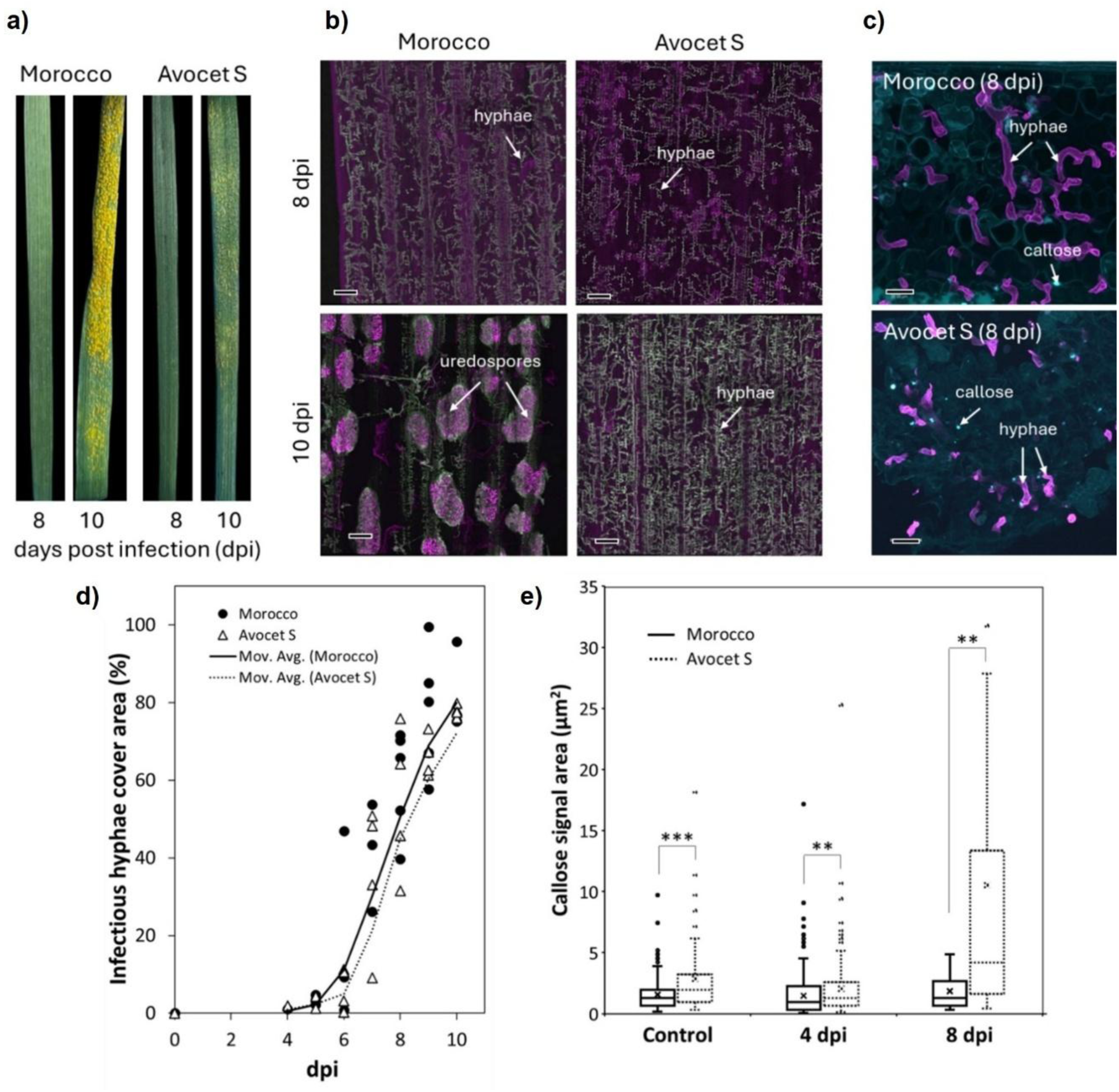
The susceptible wheat cultivar Avocet S displays a delayed infection time course to *Pst*104E137A- when compared to Morocco. a) Leaf phenotypes at 8 and 10 days post-infection (dpi). **b)** Representative confocal micrographs showing the extent of infection inside the leaves at 8 and 10 dpi. Magenta corresponds to propidium iodide-stained cell walls, and green to fungal structures stained with wheat germ agglutinin. Bars = 200 µm. **c)** Representative confocal micrographs showing callose deposition (cyan) associated with infectious hyphae (magenta). Bars = 25 µm. **d)** Infection progression and **e)** callose deposition through time in infected leaves. Asterisks denote significant differences according to Welch’s t-test at *p*-value < 0.01 (**) and *p*-value < 0.001 (***).

To assess host immune responses underlying these differences, we examined callose deposition, a hallmark of plant defence. At 8 dpi, Avocet *S* exhibited greater callose accumulation than Morocco, particularly around infection sites (Figure 6C), and this was supported by quantitative analysis (Figure 6E).

## Discussion

Recent advances in high-quality genome assemblies of *Pst* (Wang et al., 2024; Tam et al., 2025), coupled with advances in effector identification and functional genomics, have significantly enhanced our capacity to investigate host–pathogen interactions of *Pst* and other rust wheat rust fungi (Arndell et al., 2024; Chen et al., 2017; Chen et al., 2024; Cui et al., 2024; Outram et al., 2024; Salcedo et al., 2017; Shen et al., 2024; Upadhyaya et al., 2021; Wilson et al., 2024). Additionally, we have gained deeper insights into wheat defence responses by integrating comprehensive effector libraries with protoplast transient expression methods (Arndell et al., 2024; Wilson et al., 2024). Within this framework, we identified *AvrPstB48* as a hemizygous avirulence candidate effector that triggers defence signalling in a wheat cultivar-specific manner. Using wheat protoplasts, we deliberately focused on intracellular expression of *AvrPstB48,* removing the signal peptide encoding sequence from the expression clone based on localisation prediction made with EffectorP3. In cultivars such as Avocet *S, T. spelta Yr5* and Gabo, intracellular expression of *AvrPstB48* elicited strong activation of defence reporters, comparable in magnitude to the canonical *Avr/R* control *AvrSr50/Sr50*, whereas Morocco and Chinese 166 showed no detectable response to *AvrPstB48*. This clear cultivar-specific response and its intracellular recognition align with intracellular *R* gene-mediated recognition in wheat protoplasts and mirror prior observations *for AvrSr35, AvrSr27, AvrSr13, AvrSr22* and *AvrPm2a*, all of which activate the same defence reporters in protoplasts when expressed in the presence of their cognate *R* genes (Wilson et al., 2024). While we have not identified a cognate *R* gene for *AvrPstB48*, we hypothesise that it is recognised by a broadly conserved *R* gene that is common to the wheat cultivars that respond to *AvrPstB48*. Importantly, the cultivar-specific recognition of *AvrPstB48* did not correlate with full disease resistance when using the *AvrPstB48-*encoding isolate Pst104E137A– for infection assays. Several cultivars that responded to *AvrPstB48* with defence signalling in protoplasts, such as Avocet S and Gabo, were fully susceptible in whole plant infection assays at 14 dpi.

We chose Avocet *S* and Morocco for more detailed phenotypic comparisons of two fully susceptible wheat cultivars that varied in their response to *AvrPstB48*. We selected them because they are both widely used as ‘universal’ susceptible *Pst* cultivars, with the key difference that there are recent reports of *Pst* isolates that are avirulent on Avocet *S* (Nazari & El Amil, 2014) but not on Morocco. Based on this observation, there was justification to postulate that Avocet *S* carries a *R* gene designated *YrAvS* that is observable only during whole plant infection assays and with certain *Pst* isolates. We did observe a lower sporulation and hyphal coverage in Avocet *S* when compared to Morocco in our detailed macroscopic and microscopic time course infection comparison using Pst104E137A-. This infection delay in Avocet *S* is consistent with its ability to recognise *AvrPstB48,* upon which we propose the presence of the uncharacterised *YrAvS*. We do not know if *AvrPstB48* recognition contributes to infection delay and if it is recognised by *YrAvS*. An infection delay could also be caused by the recognition of another effector by *YrAvS* or any other unrelated genetic differences between Avocet *S* and Morocco. However, it is tempting to speculate that the recognition of *AvrPstB48* contributes to the infection delay, because recent segregation analysis of a *PstS8* selfing population suggests the presence of an effector that can suppress the recognition of *PstS8* on Avocet *S*, rendering *PstS8* virulent despite the presence of an avirulence effector (Livbjerg et al., 2025). Similar observations of “inhibitor” loci that suppress *Avr* recognition have been made for the flax rust fungus *Melampsora lini* (Lawrence et al., 1981) and the barley powdery mildew fungus *Blumeria graminis* f. sp. *hordei* (Caffier et al., 1996). These are consistent with a recent observation in which some *Blumeria graminis* f. sp. *tritici* isolates employ the effector SvrPm4 to suppress the recognition of AvrPm4 by Pm4 (Bernasconi et al., 2025). We therefore hypothesise that the *AvrPstB48-*Avocet *S* phenotype is explained by a ‘recognise–then–suppress’ scenario, fitting within the concept of ETSII. In ETSII, effectors that are poised to trigger *R* gene dependent disease resistance are attenuated by other effectors that suppress defence signalling, allowing disease progression despite recognition (Sohn et al., 2007; Wu & Derevnina, 2023). Therefore, the infection outcomes of the *Pst*-wheat pathosystem are likely more complex than Flor’s gene-for-gene hypothesis suggests (Flor, 1971), reflecting the complex coevolution between the obligate biotroph pathogen and its host.

Paralog analysis revealed that *AvrPstB48* belongs to a small effector family with four out of five members triggering host defence. Two paralogs (*P1* and *P2/P4*) trigger defence activation in *AvrPstB48* responsive cultivars, whereas *P3* consistently did not. *AvrPstB48_P3* provides an example of how local sequence changes within a shared scaffold mediate escape from recognition. Structural modelling and mutational analyses identified the single amino acid at position 134 as a key determinant of recognition. In AvrPstB48, P1 and P2/P4, this position encodes an aspartic acid (D134), whereas P3 encodes a glycine (G134). AlphaFold-based models predict that residue 134 lies within a solvent-exposed loop projecting from a compact, cysteine-rich core. Replacing D134 with G134 in AvrPstB48 abolished defence activation, whereas the reciprocal G134 to D134 change in P3 failed to restore recognition. This asymmetry indicates that D134 is necessary but not sufficient for recognition. The predicted *AvrPstB48* fold, a compact, cysteine-rich core likely stabilised by disulphide bonds and Zn²⁺ coordination decorated with variable surface loops, is consistent with emerging paradigms for rust effectors such as *AvrSr27*, in which conserved cysteines and metal-binding centres maintain the core, while modular loops control host specificity and escape (Outram et al., 2024).

## Acknowledgments

We acknowledge the contribution of the Plant Pathogen ‘Omics Initiative consortium in the generation of data used in this publication. The Initiative is supported by funding from Bioplatforms Australia, enabled by the Commonwealth Government National Collaborative Research Infrastructure Strategy (NCRIS).

## Author contributions

B.S., B.D., R.T., and J.P.R. conceptualized the project. B.S., R.T. and J.P.R. acquired funding. B.S. supervised the work. E.C.P., B.D., R.T., H.L., D.I.B., M.M., M.R., S.W., S.P., and F.D. acquired experimental data and conducted data analysis. E.C.P. and B.S. drafted the manuscript, and all authors contributed to writing, reviewing, and editing.

## Data availability

All supporting data is available on Zenodo (doi: 10.5281/zenodo.15845412, link: 10.5281/zenodo.15845412).

## Funding

This work was supported by a Discovery Project grant from the Australian Research Council (DP230100941) to B. S. and J. P. R. This work was supported by computational resources provided by the Australian Government through the National Computational Infrastructure (NCI) under the ANU Merit Allocation Scheme. R.T. was supported by a Grains Research and Development Corporation Graduate Research Scholarship.

## Supplementary Figures

**Figure S1.**
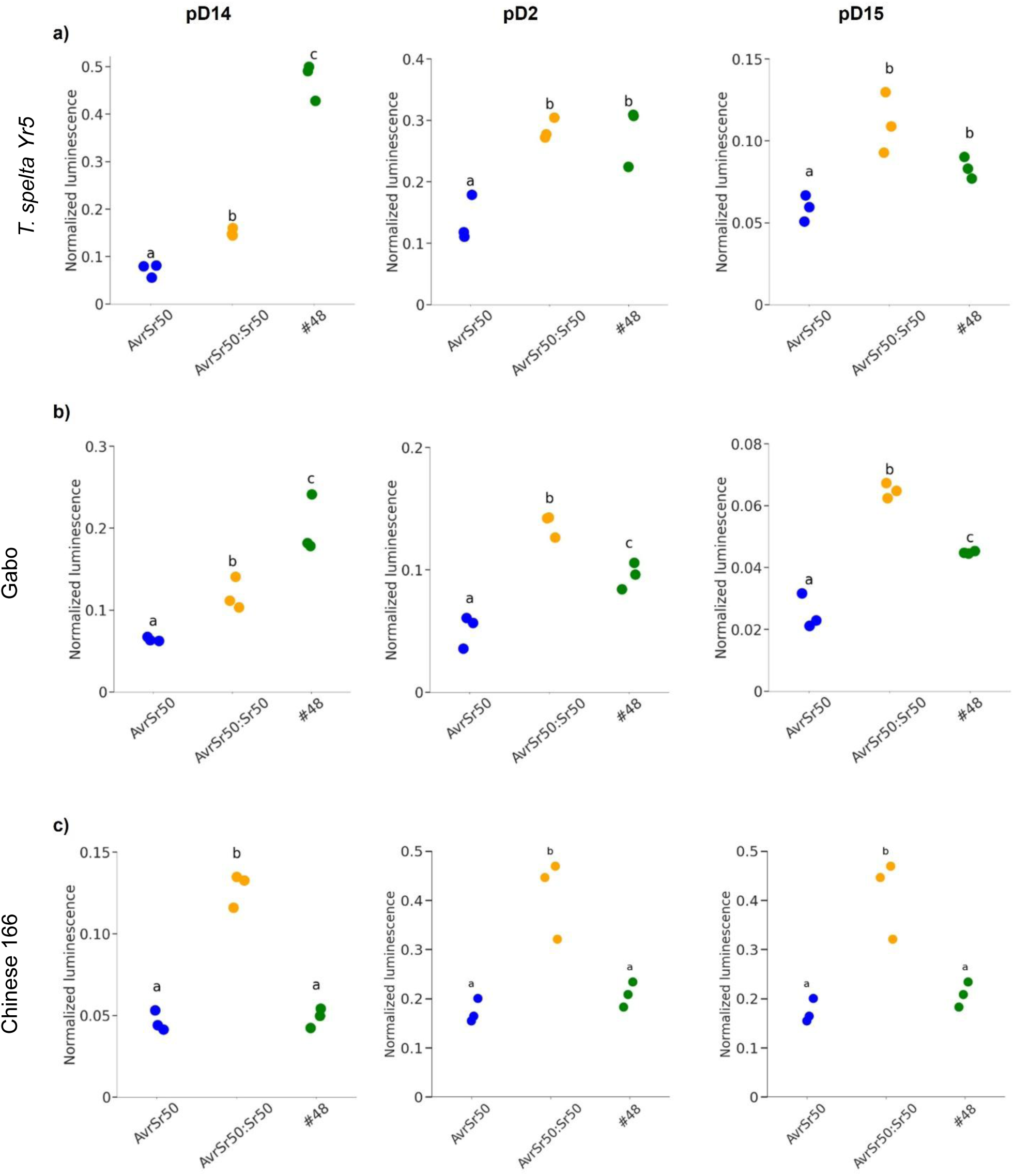
Differential activation of defence reporters by *AvrPstB48* reveals genotype-dependent recognition. Normalised luminescence signals from dual luciferase-based reporter assays using three promoter constructs (pD14, pD2, pD15) and UBI-pRedf in **a)** Triticum spelta *Yr5*, **b)** Gabo and **c)** Chinese 166. AvrSr50 and AvrSr50:Sr50 were used as negative and positive controls, respectively. One-way analysis of variance (ANOVA) followed by Tukey’s Honestly Significant Difference (HSD) test was used to compare treatments. Conditions with different letters are significantly different from each other (*p* < 0.05).

**Figure S2.**
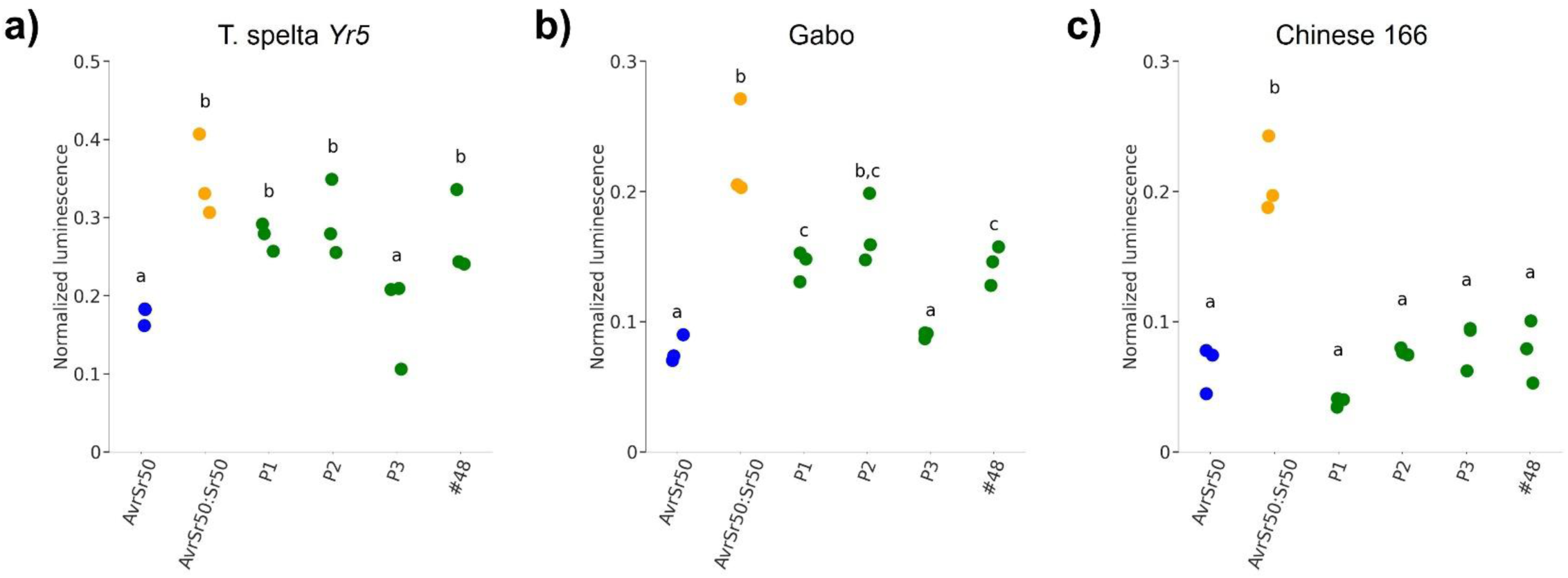
Differential activation of defence reporters by *AvrPstB48* paralogs reveals genotype-dependent recognition. Normalised luminescence signals from dual luciferase-based reporter assays using the pD14 promoter construct and UBI-pRedf in **a)** *Triticum spelta Yr5*, **b)** Gabo and **c)** Chinese 166. AvrSr50 and AvrSr50:Sr50 were included as negative and positive controls, respectively. One-way analysis of variance (ANOVA) followed by Tukey’s Honestly Significant Difference (HSD) test was used to compare treatments. Conditions with different letters are significantly different from each other (*p* < 0.05).

**Figure S3.**
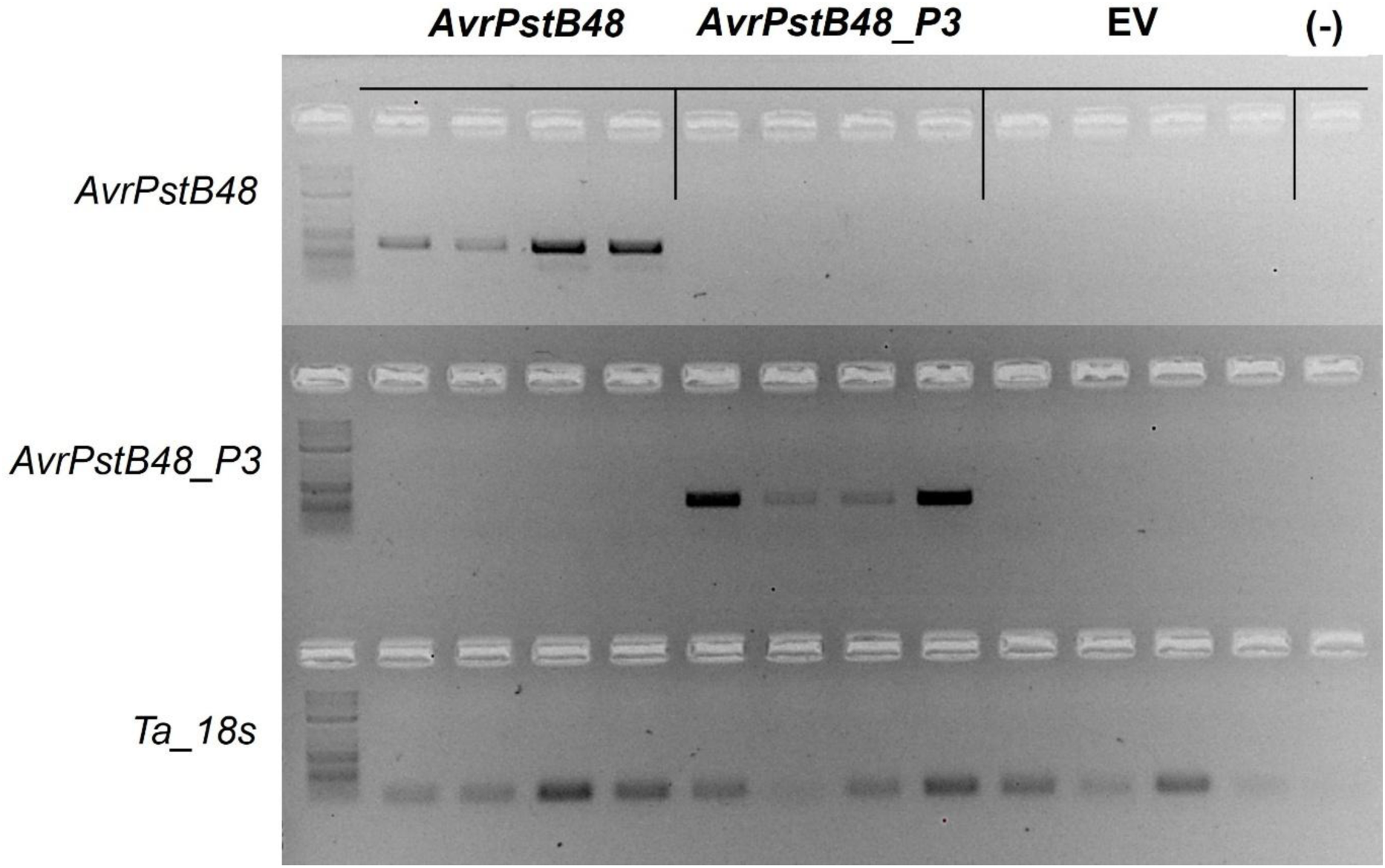
Endpoint RT-PCR confirmation of transgene expression in wheat protoplasts. Total RNA from transfected protoplasts was DNase-treated and reverse-transcribed. Gene-specific primers amplified *AvrPstB48*, *AvrPstB48_P3*, and wheat 18S rRNA (Ta 18s; loading control). Five microlitres of each amplicon were resolved on a 1% agarose gel and visualised under UV. M, 1 kb Plus DNA Ladder (NEB #N3200).

**Figure S4.**
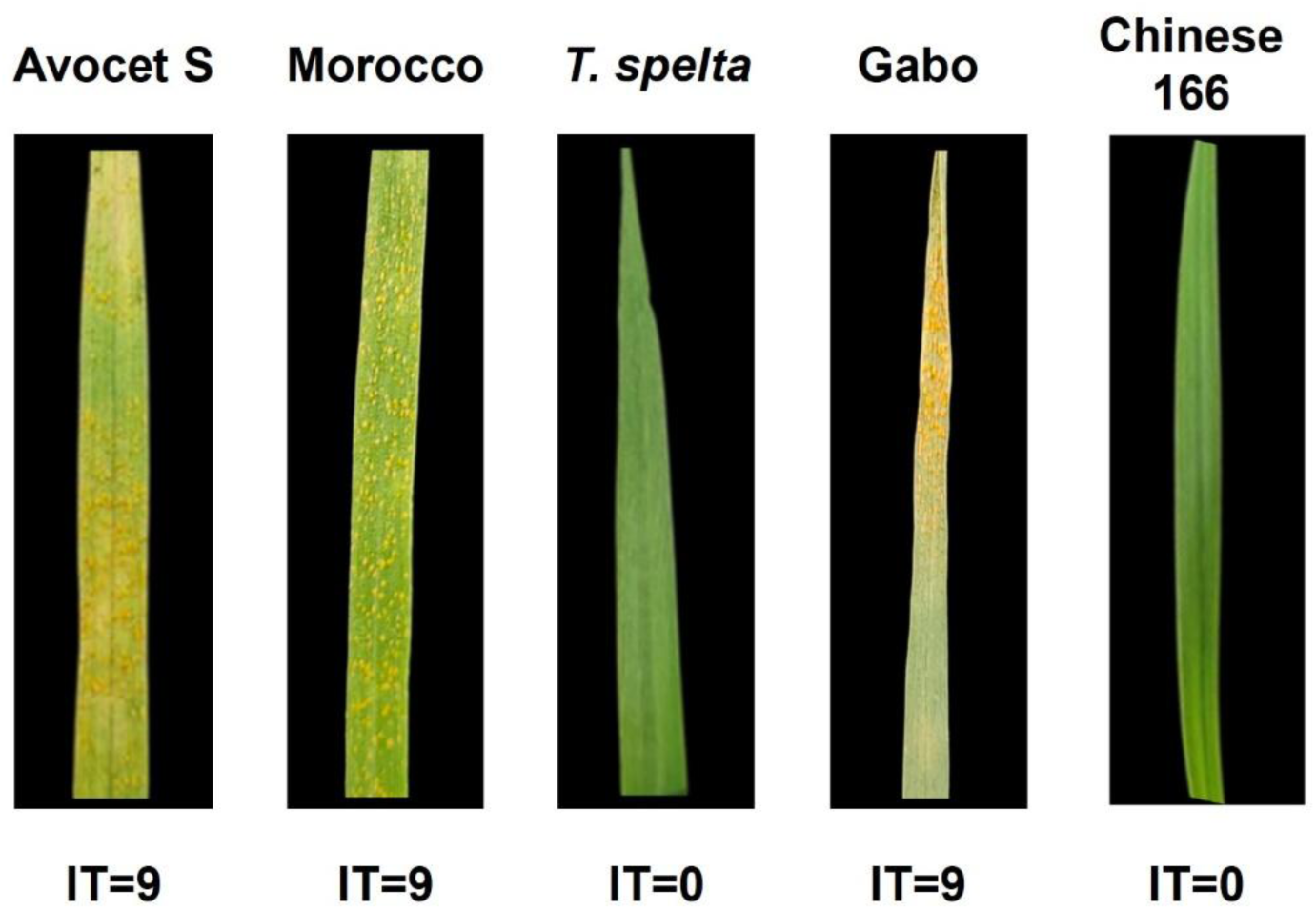
Infection types of selected wheat cultivars inoculated with Pst104E137A–. Infection types (IT) on a 0-9 scale on the wheat cultivars *Triticum spelta Yr5*, Avocet *S*, Chinese 166, Morocco and Gabo at 12 days post-infection (DPI). 0: no visible disease symptoms (immune), 1: minor chlorotic and necrotic flecks, 2: chlorotic and necrotic flecks without sporulation, 3–4: chlorotic and necrotic areas with limited sporulation, 5–6: chlorotic and necrotic areas with moderate sporulation, 7: abundant sporulation with moderate chlorosis, 8–9: abundant and dense sporulation without notable chlorosis and necrosis.

## Supplementary Tables

**Table S1.** Comparative sequence alignment and AlphaFold modelling metrics for AvrPstB48 and paralogs *(P1, P2/P4, P3*). The AlphaFold metrics include ipTM (interface predicted TM-score) and pTM (predicted TM-score), which estimate confidence in the predicted structures and inter-domain orientations, with higher values indicating greater reliability. The “Zn” column indicates whether zinc ions were included during modelling, as zinc can stabilise cysteine-rich proteins and influence predicted folding.

**Table S2.** Primers used for Avr insert verification and RT–PCR. Primer names, functions and sequences used to confirm the presence of Avr inserts (DB34, DB35) after cloning and for cDNA amplification of AvrPstB48, AvrPstB48_P3 and the wheat reference gene Ta18S in RT–PCR.

**Table S3.** List of plasmids.

**Table S4.** List of Avirulence (Avr) screening in wheat protoplasts of various differential wheat lines to *Puccinia striiformis* f. sp. *tritici* (*Pst*) Pst104E137A-infection in a protoplast-based defence reporter assay.

**Table S5.** Resume of promoter activation triggered by *AvrPstB48* and Infection types (IT) on a 0-9 scale for the isolate 104E137A- inoculated in different cultivars. Response of various differential wheat lines to *Puccinia striiformis* f. sp. *tritici* (*Pst*) *Pst*104E137A-infection and the induction of *AvrPstB48* in a protoplast-based defence reporter assay. “*Pst*-resistant gene” refers to the known Yr gene(s) present in each line. “*AvrPstB48* induction” indicates whether candidate *AvrPstB48* triggered a defence response (+) or not (-) in the protoplast assay. “*Pst* infection *outcomes*” represents the infection types (IT) on a 0-9 scale on the wheat cultivars at 12 days post-infection (DPI). 0: no visible disease symptoms (immune), 1: minor chlorotic and necrotic flecks, 2: chlorotic and necrotic flecks without sporulation, 3–4: chlorotic and necrotic areas with limited sporulation, 5–6: chlorotic and necrotic areas with moderate sporulation, 7: abundant sporulation with moderate chlorosis, 8–9: abundant and dense sporulation without notable chlorosis and necrosis.

## Notes

### Competing Interest Statement

The authors have declared no competing interest.

### Summary of Updates

In this revised version, we have strengthened the manuscript by adding new data and clarifying our approach to effector expression. We have also substantially revised the text to improve clarity and accuracy. The Results and Discussion sections were restructured to provide a more data-driven interpretation of the Avocet S versus Morocco phenotypes, to clarify the proposed recognise then suppress (ETS II) model, and to refine the description of effector family structure and loop epistasis. Several figures and table legends were improved to aid interpretation.

